# Improved spatio-temporal measurements of visually-evoked fields using optically-pumped magnetometers

**DOI:** 10.1101/2021.01.27.428396

**Authors:** Aikaterini Gialopsou, Christopher Abel, Timothy M. James, Thomas Coussens, Mark G. Bason, Reuben Puddy, Francesco Di Lorenzo, Katharina Rolfs, Jens Voigt, Tilmann Sander, Mara Cercignani, Peter Krüger

## Abstract

Recent developments in performance and practicality of optically pumped magnetometers have enabled new capabilities in non-invasive brain function mapping through magnetoencephalography. In particular the lack of need of cryogenic operating conditions allows for more flexible placement of the sensor heads closer to the brain surface, leading to improved spatial measurement resolution and increased source localisation capabilities. Through the recording of visually evoked brain fields (VEF), we demonstrate that the closer sensor proximity can be further exploited to improve the temporal resolution. We use optically pumped magnetometers (OPMs), and for reference superconducting quantum interference devices (SQUIDs), to measure brain responses to standard flash and pattern reversal stimuli. We find highly reproducible signals with consistency across multiple healthy participants, stimulus paradigms and sensor modalities. The temporal resolution advantage of OPMs is manifest in a fourfold improvement of the ratio of magnetic signal peak height to temporal width, compared to SQUIDs. The resulting capability of improved spatio-temporal signal tracing is illustrated by simultaneous vector recordings of VEFs in the primary (V1) and associative (V2) visual cortex, where a time lag on the order of 10-20 ms is consistently found. This paves the way for further studies of spatio-temporal neurophysiological signal tracking in visual stimulus processing and other brain responses with potentially far-reaching consequences for time-critical mapping of functionality in the healthy and pathological brains.

## 1. Introduction

Over the last century, outstanding advances in medical physics have led to the development of non-invasive functional neuroimaging techniques [1–3]. This has provided significant insights into brain function and connectivity. Important improvements in modern neuroimaging techniques have allowed neural patterns associated with specific stimulations to be investigated [4], providing information about the signal’s spatial and temporal characteristics [5]. Previous studies have shown that a spatio-temporal analysis of brain signals is not only essential to understand the basic mechanisms of brain circuits, but would also provide reliable biomarkers for differentiating physiological and pathological brain activity in neurodegenerative diseases [6, 7]. There is even a potential for predicting clinical progression or treatment responses [8]. The realisation of the full scope of temporal and spatial localisation of brain signals, however, is hampered by the intrinsically low spatio-temporal resolution of currently available methods [9, 10].

Functional Magnetic Resonance Imaging is capable of mapping activated brain regions with high spatial resolution, but offers only low temporal resolutions (~1 s), as the local measured changes in blood flow are not synchronized with neuronal activity [11]. Electroencephalography (EEG) is a real-time neuroimaging method, with limited source localisation capability and spatial resolution (~10 mm) [12].

Magnetoencephalography (MEG) is an alternative real-time method with a theoretically possible improved spatial resolution, able to measure postsynaptic potentials of tangential pyramidal cells at the surface of the scalp [12]. Recent research has shown that MEG can be used for the evaluation of abnormal cortical signals in patients with Alzheimer’s disease [13], Parkinson’s disease [14], autism spectrum disorder [15], and in severe cases of post-traumatic stress disorder [16]. However, MEG suffers from low signal-to-noise ratio (SnR), and its use is confined to magnetically-shielded rooms (MSRs). The magnetically shielded environments are used to subdue environmental magnetic noise, often many orders of magnitude higher than neuromagnetic fields (fT to pT range).

Traditionally, MEG relies on an array of super-conductive quantum interference devices (SQUIDs) to measure the brain’s magnetic fields [17].

With the sensor array being fixed inside a required cryogenic dewar, the locations of the individual sensors must be arranged to fit a vast majority of head sizes and shapes [18]. The fixed positions result in different radial offsets from a subject’s head. Coupled with tiny head movements from a subject during a measurement, the offsets and fixed positions have a major impact on the potential cortical activity detection [19]. In particular, the theoretically achievable precision of signal source localisation is lost. This makes SQUID-MEG impractical in many cases, in particular in the clinical context.

Extremely sensitive spin-exchange relaxation-free (SERF) optically-pumped magnetometers (OPMs), developed at the turn of the millennium [20], can help to overcome the SQUID-MEG limited spatial resolution [21]. With the OPMs able to be fixed to a subject’s head [22], a smaller offset distance than SQUIDS, and the ability for simultaneous dual axis measurements, OPM-MEG has several advantages over SQUID-MEG, including its suitability for applications within pediatric and clinical populations.

The aim of this study was to demonstrate the improved ability of OPM-MEG by recording spatio-temporal characteristics of neurophysiological signals, and comparing them to conventional SQUID-MEG. As a prototypical test case we have chosen visual cortex responses to established standard visual stimulations, with the measured responses evaluated in a well-characterised context. We find that OPM-MEG is superior to SQUID-MEG in brain signal tracking in space and time, making a suitable method to provide new information about propagating signals, source localisation, neural speed, and brain circuits far beyond the processing of visual stimuli.

## 2. THEORY CONSIDERATIONS

### 2.1. Temporal resolution

A typical response to a given brain stimulation is measured in MEG as a time-sequence of a series of peaks. The peak timing is important for the understanding of key brain functions. It is also important to accurately determine the time of the peak for comparison between different sensors and participants.

Due to large variations in individual signals, MEG signals are commonly averaged over many trials. This averaging over uncorrelated measurements enables association of a statistical standard error with the mean signal determined at each point in time. We define the timing error on a peak to be given by the time range in which the peak signal value remains within the standard error. This timing error is often larger than the sample rate.

In practice, the sample rate is hence not typically the limiting factor in determining the timing of characteristic signal patterns. We argue instead that more accurate timing, i.e. improved temporal resolution, can be obtained when the ratio between signal strength and temporal width of a feature (or peak *height-to-width ratio* HWR) is maximised. For a quantitative demonstration, we use a simple model of a peak with a Gaussian shape.

The uncertainty in the time at which the peak occurs can be calculated by,

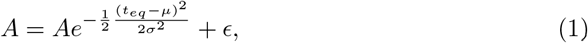

where *A* is the amplitude of the peak, *t_eq_* is the time that the peak and the error upper bound have the same value, *μ* is the time of the peak, 2*σ*^2^ is the width of the Gaussian distribution and *ϵ* is the standard error, assumed to be constant in this example. We can solve for *t_eq_* finding,

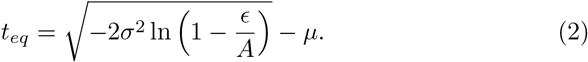

Note this only holds with *ϵ*/*A* < 1. In first order Taylor expansion (in the logarithmic term) this simplifies to

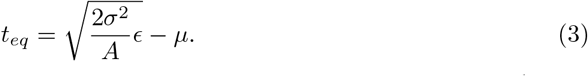

The error on the peak position is 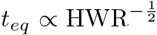 with HWR = *A*/2*σ*^2^. Whilst this derivation has been shown for a Gaussian, a similar argument can be made for any peak shape.

### 2.2. Spatial resolution

As the magnetic field amplitude decays according to a power law with the distance from a field source, improved signal detection is achieved when sensors are moved closer to the brain. As shown formally for a generic situation [23], and confirmed through realistic brain anatomy simulations [24, 25], the consequences of the field decay law are that closer positioning of a sensor system provides improved signal-to-noise, better spatial resolution and more precise source localisation. In general, when applying the Rayleigh criterion for resolution, the maximum distance at which two sources can be resolved is comparable to the distance between the two sources [23]. As OPMs can be placed closer to the head than SQUID systems, the OPMs are able to achieve a higher spatial resolution.

### 2.3. Vectoral measurements

In conventional MEG only one component of the vectorial magnetic field is measured. Most commercial setups for SQUID-MEG only measure magnetic field gradients radial to the brain. At the typical stand-off distances of several centimetres, the orthogonal components tend to be weak, so that the radial field (gradient) component approximates the total field (gradient). With closer sensor proximity to the brain, OPMs are able to measure multiple field (gradient) components to extract additional spatial information [26]. A vector measurement taken at short distances does not suffer from the zone of a vanishing field component in the immediate vicinity of a current dipole, and is sensitive to volume currents in the brain.

Measuring both radial and tangential field components also helps to improve signal *temporal* resolution. This is a consequence of the ability to characterise the field as a vector. At the sensor, the magnetic field has a direction and magnitude. A radial sensor measures the magnetic field projected onto the radial direction. By measuring in only the radial direction it is not possible to differentiate between a rotation or a change in magnitude of the magnetic field vector. Worse still, if the magnetic field vector simultaneously changes in both direction and magnitude, then the time at which the magnetic field reaches peak magnitude can be obscured. By measuring a second component of the magnetic field we can begin to differentiate between a change in the magnitude of the magnetic field, and a change in magnetic field direction. Sensors near the head are in a source-free region, therefore using Ampère’s Law ∇ × *B* = 0 the third magnetic field component can be calculated from the other two magnetic field components assuming the gradient of the magnetic field can be calculated. For a system with a low sensor count, all three magnetic field components need to be measured to achieve full characterisation of the magnetic field.

## 3. MATERIALS AND METHODS

### 3.1. Participants and MRI

Visual evoked fields were studied in 3 healthy participants (2 men aged 26 and 30, 1 woman aged 47 years) with normal or corrected-to-normal vision. The 3 participants received a 3 T MRI scan (Siemens Magnetom Prisma, Siemens Healthineers, Erlangen, Germany) at the University of Sussex, including a high-resolution T1-weighted anatomical scan. For one participant a diffusion-weighted scan was acquired for reconstructing the optic radiations, with two diffusion-weighting shells (b values = 1000 and 3000 s/mm^2^). For each b value, diffusion gradients were applied along 60 non-collinear directions. Six images with no diffusion weighting (b=0) were also collected. Image processing was performed using tools from the FMRIB’s Diffusion Toolbox 5.0. First, data were corrected for involuntary motion and eddy currents using affine registration. BEDPOSTx was run with default settings to fit a crossing fibers model [27], and finally, XTRACT was used to automatically reconstruct the left and right optic radiations in native space by probabilistic tractography [28]. The results are shown in Fig. 1(d) and (f).

**Figure 1.**
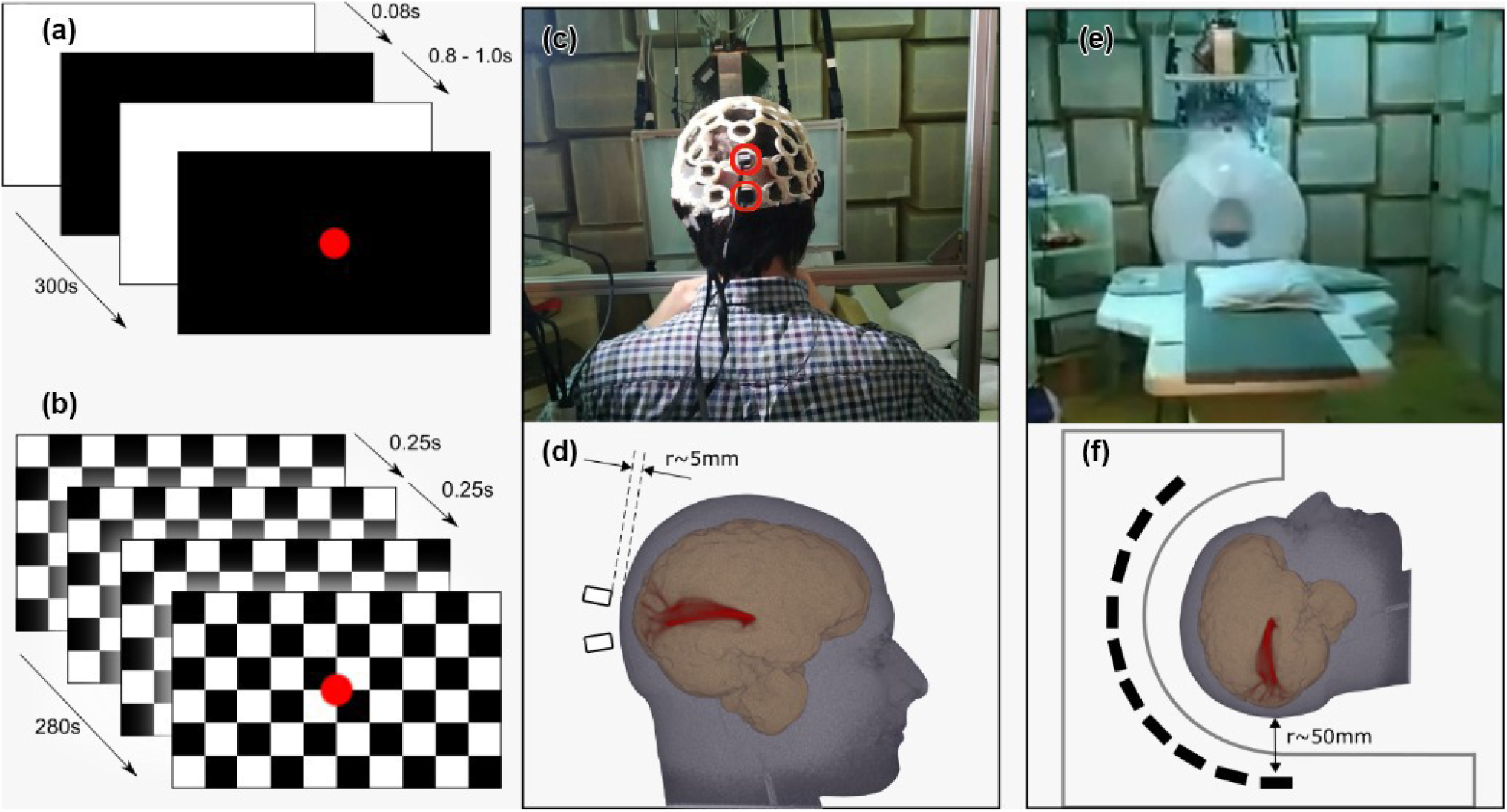
(a) Flash and (b) pattern reversal stimulation protocols. (c) A participant in position with the 3D printed helmet containing the OPM devices. Red highlighted cells show the sensor locations used for the study. (d) 3D rendering of the MRI scan of Participant 1 showing approximate locations of OPM sensors 1 & 2, and scalp-sensor separation of around 5 mm. The reconstructed optic radiation are also shown in red. (e) The Yokogawa SQUID-MEG system. (f) Schematic of the SQUID-MEG system showing a sensor to scalp separation of approximately 50 mm.

### 3.2. Experimental Design

The study was approved by the Brighton and Sussex Medical School Research Governance and Ethics Committee (ER/BSMS3100/1), and all participants gave written informed consent to take part, after explanation of the procedure and purpose of the experiment. All MEG measurements, OPM-MEG and SQUID-MEG, were taken in the Ak3b MSR (Vacuumschmelze, Hanau, Germany) at Physikalisch-Technische Bundesanstalt (PTB), Berlin. The MSR is equipped with an external triaxial active shielding coil system controlled by fluxgates. Inside the MSR field fluctuations are sufficiently weak to allow OPM operation [29, 30].

Two standard full-field visual stimulation protocols were employed during the MEG recording, a flash stimulus (FS), and a pattern reversal stimulus (PR). The parameters used were based on standard guidelines for clinically evoked potentials [31]. These paradigms are widely used to evaluate early visual processing, and to detect abnormalities in the visual pathways. The flash stimulus, shown in Fig. 1(a), consists of short white flashes of length 0.08 s (5 frames). To avoid participants from preempting the stimulus, each white flash was followed by a dark period with the length varying pseudo-randomly between 0.92 s and 1.00 s (55 to 60 frames). The total duration of a single FS measurement run was 300 s.

The pattern-reversal stimulus, figure 1(b), consisted of a black and white checkerboard (10 squares wide, 8 high) with the colours inverting at 0.5 s (30 frame) intervals. Each run had a duration of 280 s. For both FS and PR, a red dot was continuously projected onto the centre of the screen to act as a focal point for the participant. Before each measurement run, whilst in position for the trial, the participants were exposed to a three-minute dark adaptation period. Measurements of the empty MSR were obtained in order to evaluate environmental noise levels. During the noise measurements, the OPMs were located in the same position and orientation as they would with a participant wearing the sensors. During measurements, participants sat upright with the sensors mounted in a 3D-printed helmet (figure 1(c)). A chin rest was used to help stabilise each subject’s head, reducing movement when looking forward at a 50 cm × 34 cm vertically orientated screen. The stimuli were projected on to the screen via a mirror system using a 60 Hz LCD projector, positioned outside of the MSR. The SQUID-MEG system (figure 1(e)) accommodated participants in a horizontal position, with the same screen positioned horizontally above the subject. The screen to eye distance was 53 cm for the OPM-MEG setup and 45 cm for the SQUID-MEG system.

## 4. MEG SYSTEMS OVERVIEW

### 4.1. OPM-MEG

The OPM-MEG system consisted of two second-generation QuSpin zero-field magnetometers (QuSpin Inc., Louisville, CO, USA), with a specified typical sensitivity <15 fT/Hz^1/2^ and magnetic field measurement bandwidth of 135 Hz in a 12.4 mm × 16.6 mm × 24.4 mm sensor head. The OPMs were mounted in a 3D printed helmet (open-source design; OpenBCI Mark IV helmet) ‡ and positioned over the visual cortex at the Oz and POz positions, according to the standard 10-10 system [32]. These locations were chosen in accordance with each subject’s MRI scan.

The Oz and POz positions correspond to the primary visual cortex (V1) and the associative visual cortex (V2), respectively. Studies have shown the feed-forward and feedback interaction between the V1 and V2 areas in response to visual stimulation [33]. More specifically, there is an early activation at V1, known as the P1 or C1 component, which is then suppressed as the signal propagates to V2, after which a reflected wave is initiated and propagates back to V1 [34].

The design of the helmet and sensor head fixes the scalp to sensor distance to ~5 mm. Python-based software was developed for the design and presentation of stimuli. The software was directly connected and synchronized with a main OPM-MEG data acquisition system (DAQ). The OPM-MEG system’s analogue outputs were recorded at 1 kHz via a Labjack T7 pro (Labjack Corporation, Co, USA). All DAQ electronics, except the OPM sensor-heads, were located outside the MSR and directly connected to the Labjack and control computer.

### 4.2. SQUID-MEG

The SQUID-MEG system MEGvision (Yokogawa Electric Corporation, Japan) comprises of 125 axial gradiometers and 3 reference magnetometers. For the stimuli presentation the same software was used to prevent any bias in the stimulation delivery. Data from all sensors were recorded at a 2 kHz sampling frequency, and the sensors located closest to the OPM positions were used for analysis. The fixed positions of the sensors result in a ~50 mm standoff from the subject’s scalp. The SQUID-MEG used MEG Laboratory 2.004C (Eagle Technology Corporation) data acquisition software.

### 4.3. Data Analysis

The DAQ systems were synchronised with the presentation software. Both OPM and SQUID data analyses were performed using the FieldTrip toolbox [35] and MATLAB. In order to isolate the frequencies of interest with relevance in visually evoked fields, all data were filtered with a bandpass filter between 5 and 60 Hz. A further bandstop filter was applied between 49 and 51 Hz to suppress 50 Hz line-noise. The epoched trials for FS were −45 ms to 350 ms, and 0 ms to 250 ms for PR. Any trials with interrupted recordings were removed from the analysis. All the time-locked averaged responses contain more than 380 trials for the FS stimulation, and more than 280 for PR.

In the following sections the evoked fields are shown as the mean across all individual trials for a single run. The uncertainties on the signal amplitudes are calculated as the standard error at each time point (with a 1 ms time spacing for OPM-MEG and 0.5 ms for SQUID-MEG). The resulting uncertainty band is then used to determine temporal uncertainties of signal features such as amplitude peaks. The time error is set as the width of the uncertainty band at the amplitude feature, as outlined in Sec. 2.1.

In order to compare the spatio-temporal response of OPM-MEG to SQUID-MEG we initially study the temporal resolution of the two systems by measuring the signal height to (temporal) width ratio (HWR) in characteristic evoked magnetic field peaks. In each visually evoked field (VEF) we found the dominant peaks for each sensor type and estimated the HWR. The signal height is taken as the difference between the signal at the peak maximum and the mean of the two adjacent local minima. The width is set as the time difference between the two local minima (see insert in Figure 2). The HWR uncertainty results as error propagated from time and signal uncertainties, determined in the above described manner.

**Figure 2.**
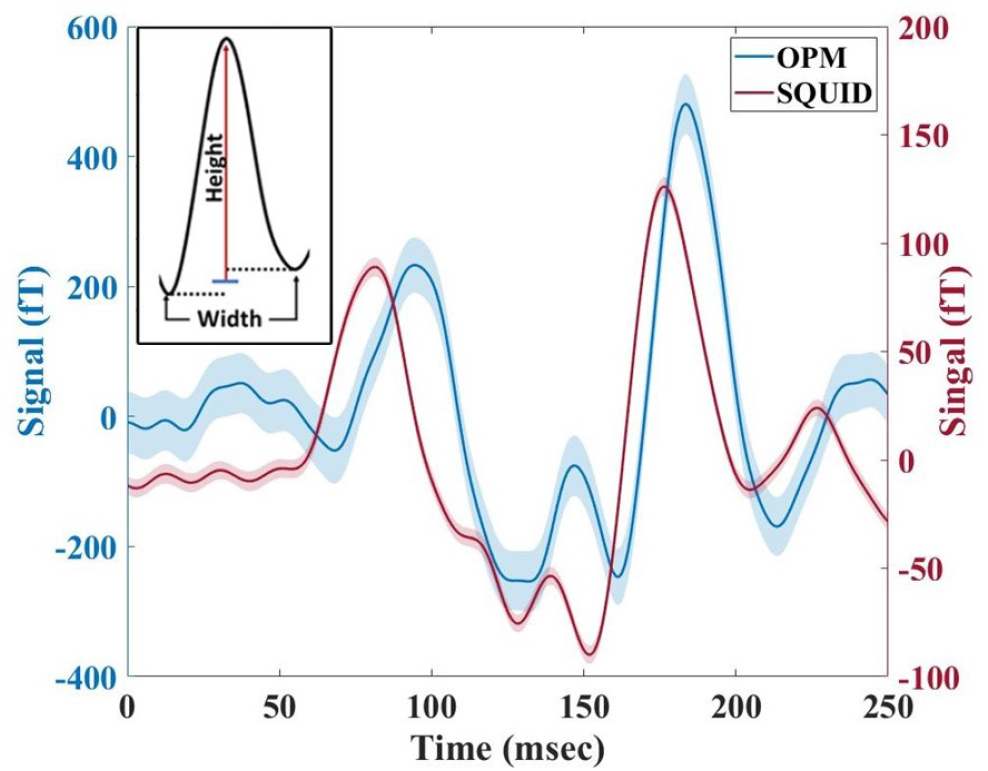
Averaged evoked field recorded by OPM-MEG and SQUID-MEG for Participant 1 for flash stimulation over a single measurement run (300 trials). VEF measured at Oz using OPM sensor (blue line) and the corresponding SQUID sensor (red line).The shaded area shows the standard error. Inset: The signal height (red line) is the amplitude difference between the peak maximum and the mean of the two local minima (blue line). The width is the time difference between the two local minima (dashed lines). The HWR is the ratio of these two values.

In a second step, the evoked potentials as measured at the two sensor locations are then compared. For the OPM system, the simultaneously obtained individual field component (radial *B_z_* and axial *B_y_*) data are further compared to the resulting planar projection *B_yz_*, with 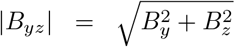. This reduces timing artefacts that can occur in data restricted to a single component.

VEFs are characterized by three time components occurring at different times: the early component (P1), the main component (P2 for flash stimuli and P100 for pattern reversal stimuli), and the late component (P3), where we established the onset range for the main and late components based on previous studies [10, 36–40]. The flash and pattern reversal stimulus responses consist of an early component with peak onset between 35 and 60 ms, a main component (P2) between 83 and 152 ms, and a late component (P3) between 160 and 230 ms.

Each participant had at least four FS runs and three PR runs with the OPM-MEG system, and a single run for each stimuli with the SQUID-MEG.

The averaged responses were determined using the same method as detailed above.

## 5. RESULTS

In total, 30 OPM-MEG runs were undertaken, with 12 FS, 9 PR, and 9 background runs. In addition, a single FS and PR run was conducted with SQUID-MEG for each participant. The VEFs from all participants and all modalities were consistent with patterns known from the literature.

We consider the HWR of the two systems and the higher SnR of OPM-MEG observed in other studies [22, 24]. Figure 2 shows measurements from a single OPM sensor and the corresponding SQUID sensor for FS. The OPM-MEG recorded signals with up to 4 times higher amplitude than the SQUID-MEG, with the OPM and SQUID sensors recording a maximum amplitude of 480(46) fT and 126(4) fT, respectively. Along with the increase in amplitude over the SQUIDs, we see the same activation patterns in both methods, further verifying the OPM’s recorded traces. The OPM HWR was found to be 13.4(2) fT/ms, compared to the SQUID HWR of 3.4(2) fT/ms. In spite of us observing a factor ~2.5 improvement of the SQUID SnR over the OPM-MEG, in our case the higher HWR still indicates a higher temporal resolution of the OPM-MEG neuroimaging system. The OPM VEF shows more pronounced peaks, resulting in sharply defined maxima and minima, resulting in lower temporal uncertainties compared to SQUID-MEG measurements.

In figure 3 we plot all participant 1 PR and FS runs recorded by OPM-MEG at Oz, along with their average. The individual runs illustrate the reproducibility of the activation patterns during both stimulations, with the main (P2 or P100) and late components (P3) having similar time onsets across all runs. For FS, the main component (P2) has an onset time between 135 ms and 100 ms and the late component (P3) between 180 ms and 190 ms. For PR, the main component P100 occurs between 128 ms and 133 ms, while the late component (P3) occurs between 210 ms and 214 ms.

**Figure 3.**
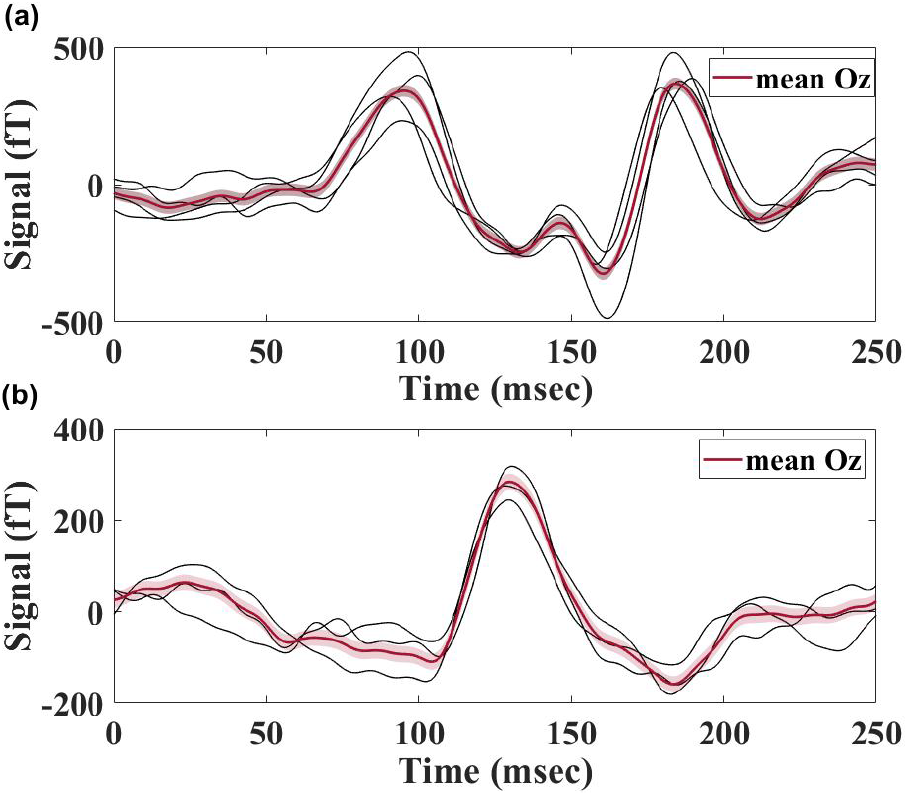
OPM VEF for (a) four FS runs and the associated mean. (b) Three PR runs along with the mean for Participant 1. The individual runs (black) for both FS and PR show the same activation pattern as the associated mean (red). The shaded area displays the standard error of the mean.

In order to quantify the within-participant reproducibility of VEFs qualitatively observed in Fig 3, we calculate the Pearson correlation coefficients between runs per participant. The correlation coefficients for the flash stimulus and pattern reversal were calculated to be 0.83(7) and 0.85(3) for participant 1, respectively, and 0.70(13) and 0.14(27) for participant 2, and 0.65(21) and 0.54(9) for participant 3. In addition to this, we then calculate the correlation coefficient between subjects for the same stimuli. Moderate between-subject correlation coefficients were found for participants 2 and 3 (0.38 for Oz), while Participant 1 showed anti-correlated signal at both sensors (−0.53 with participant 2;−0.54 with participant 3). The different cortical folding of each participant could explain the anti-correlation measured between the VEFs. Previous studies have shown an asymmetry in extracranial magnetic field measurements due to variabilities in cortical folding [41, 42].

Figure 4 displays a single run recorded by OPM-MEG (a) and SQUID-MEG (b) systems during FS. Although OPM-MEG shows the initial activation (P1) at the primary visual cortex, there is a significant time difference between the arrival of the signal at POz and Oz for both the main and late components, with an earlier activation at POz. The vertical purple bands represent the time range of P2 and P3 components found in previous studies for Oz EEG sensors [9, 10, 36–40]. The dominant peaks that fall within these boundaries are shown by bold dashed lines, representing the peak times of the main (P2) and late (P3) components. Δ*τ*_1_ and Δ*τ*_2_, defined as the delay between signals arriving at POz and Oz for the main and late components, were measured as 10(7) ms and 20(4) ms, respectively. The earlier activation of POz compared to Oz for Participant 1 was observed in all four runs, with 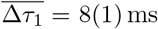 and 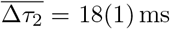 (see Fig. 5).

**Figure 4.**
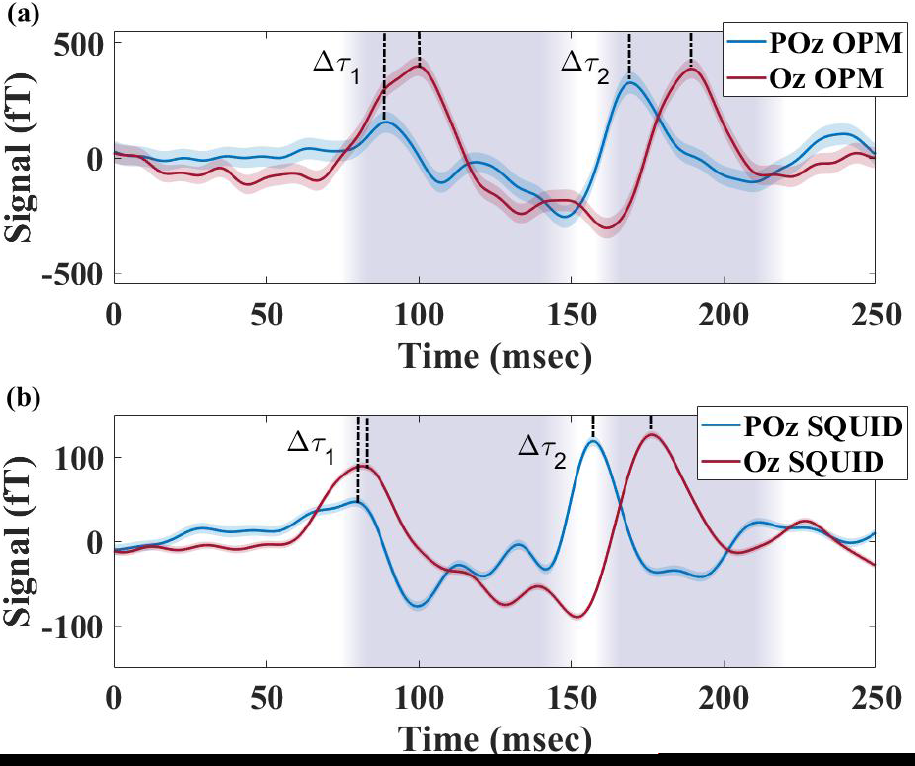
Visually evoked response during flash stimulation recorded by: a) OPM-MEG and b)SQUID-MEG for Participant 1. The coloured areas indicate the limits where the peak onset for Oz is expected for each stimulus [10, 36–40]. The selected peaks for Oz (red) and POz (blue) sensors are marked with dashed lines for both components P2 and P3.

**Figure 5.**
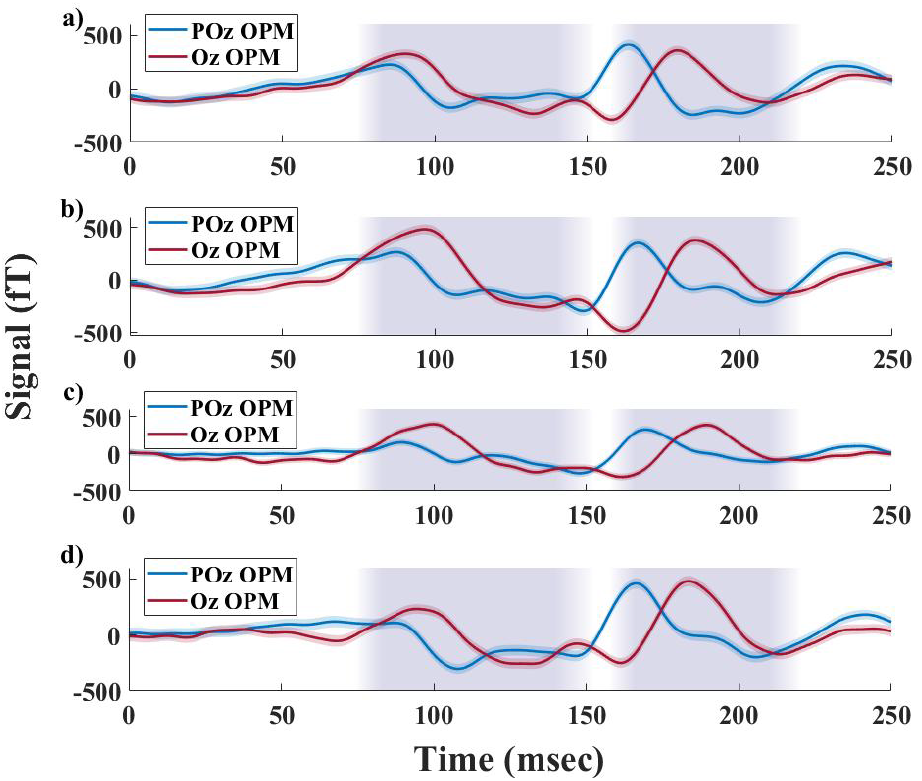
OPM VEF between the POz and Oz sensor for four flash stimulation runs for Participant 1. All four runs show similar peak onset and amplitude. The time lag between the POz (blue) and the Oz (red) OPM sensors is consistent for the two components across the runs.

The reproducibility of the time delay, and the small variations in Δ*τ*_1_ and Δ*τ*_2_ over multiple measurement runs points to a neurophysiological origin of the delay, such as a timing difference of signals arriving at different locations within the visual cortex. Similar time delays are also suggested in Figure 4(b) for the SQUID-MEG measurement, where we find Δ*τ*_1_ = 2(5) ms and Δ*τ*_2_ = 18(3) ms. Although we recorded a higher SnR for SQUID, over OPM measurements, the timing uncertainties for Δ*τ*_1,2_ are similar in both modalities due the improved HWR achieved with OPMs. The observed activation patterns were shown to be reproducible across all runs (Fig. 5) and stimuli (Fig. 4), with activation of POz before Oz detected in all participants.

Compared to SQUIDs, the additional feature of OPM-MEG to simultaneously measure components along two axes, in this case *y* and *z*, can be used to further support the neurophysiological origins of the delay phenomenon, such as those observed here for the signal’s main and late components. Figure 6 shows the two magnetic field components *B_y_* and *B_z_* simultaneously measured by two OPMs, along with each sensor’s magnitude projected in the *yz*-plane |*B_yz_*|. In Figure 6(a) we show OPM-MEG *B_y_* and *B_z_* FS responses recorded simultaneously from a single run at POz and Oz. *B_z_* shows a VEF with higher amplitudes and more clearly discernable peak structure than that recorded by *B_y_*. In Figure 6(b) we show |*B_yz_*|. The characteristic components of the VEF recorded in the vector components persists, including the timings and relative time delays of the main VEF features (previously negative peaks are now positive as the *yz*-plane projection is displayed as the modulus). Our result of a significant and reproducible time delay between signals arriving at POz and Oz (Figures 4 and 6) is consistently observed across participants and stimuli.

**Figure 6.**
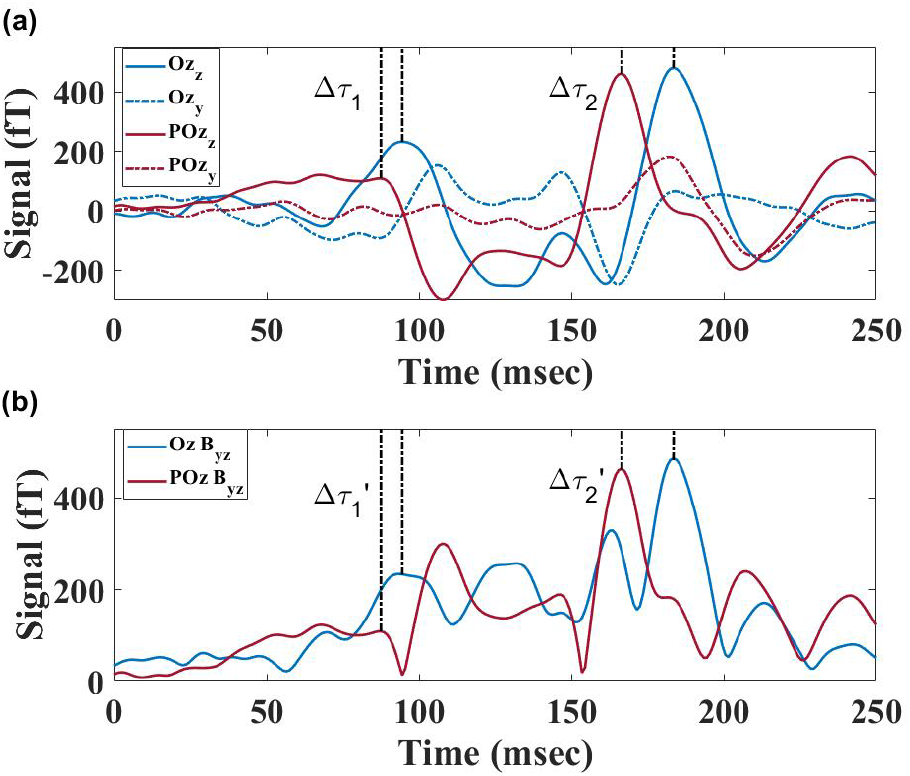
FS VEF recorded at Oz (blue) and POz (red) using OPM-MEG. a) The averaged Oz and POz response along the *z* (bold) and *y* (dashed) directions. b) The Oz and POz magnitude projected into the y-z plane. The bold black lines indicate the earlier activation of POz followed by the Oz.

## 6. DISCUSSION

In this study we used VEFs to assess and demonstrate the ability of MEG based on two types of highly sensitive magnetometers (OPMs and SQUIDs) to detect neurophysiogical brain signals with simultaneously high spatial and temporal resolution. We find that both sensor modalities are suitable to reproduce characteristic brain signatures known from well-established neurophysiological research and clinical practice.

The ability to track local brain responses in space and time can be quantified by determining the time interval over which a signal rises and falls, i.e. the ratio between the amplitude of a peak signal and the temporal width of the peak. We find that the OPM HWR has a fourfold increase over SQUID measurements, confirming the expectation of the closer proximity of the OPMs to the visual cortex having such an effect.

Importantly, we were able to confirm that the OPM-MEG measurements are robust. Repeating the experiment with two different visual stimuli (flash stimulus and a checker board pattern reversal stimulus) and with three different participants, we observed good reproducibility over multiple repeated runs within each subject and each stimulus. Differences between subjects and type of stimulus are discernable, but the key signal characteristics remain. Individual cortical folding variations could lead to different cancellation of the extracranial magnetic field [41, 42] which reflects the asymmetry of the VEF and the anti-correlation between some of the participants’ responses.

Finally, we illustrate that OPM-MEG is capable of recording neurophysiological signals of a common origin at different locations at different times. By measuring the arrival times of characteristic VEFs at two distinct locations within the visual cortex. The temporal resolution is sufficiently high to determine significant time differences between the primary visual cortex (Oz) and the associative visual cortex (POz). We measured a consistent delayed response at the Oz position relative to the POz position on the order of 10 to 20 ms for the second (P2/P100) and third (P3) components. This observation is again highly reproducible for different runs and is similar across participants and both types of stimuli. It is confirmed by corresponding SQUID measurements. The time delay uncertainties of the OPM data are comparable and even slightly lower than their SQUID counterparts. We attribute this to the strongly enhanced HWR featured by the OPMs.

In order to verify the neuronal nature of the measured time delay, we analysed the recordings along the OPM’s orthogonal *y* and *z* axes. Although we have already demonstrated that our analysis of *B_z_* results in an earlier activation of POz, the inclusion of a second axis, for which we now measure |*B_yz_*|, follows the same trend. We can indicate that the observed activation pattern is more likely to be from neural activity than an artifact of limited information. The future addition of a third orthogonal axis, to complement our two axis system, will be beneficial in order to fully validate the activation source observed. As we have demonstrated the reproducibility of VEFs in separate runs, sometimes recorded over different days, acquiring three dimensional recordings by rotating the OPM-MEG sensors between runs could be used in future experiments.

Although the VEFs are well defined in humans [39], the spatio-temporal pattern of the propagating signal is not well characterized. Studies have previously revealed the interaction of the primary visual cortex (V1) with associative visual areas (V2, V3) using an invasive cortical feedback system in animal models [33, 34]. The hierarchical order and spatio-temporal processing of the signal in humans remains uncertain. Some studies have claimed P1 originates from the primary visual cortex [10, 43], while others indicate it originates from the extrastriate cortex [36, 44]. Additionally, the P2/P100 component appears to originate from the extrastriate cortex without a definite region [45]. The widespread sensor positioning of electrodes or SQUIDs combined with the low spatio-temporal resolution may not be able to record coincident responses from close cortical sources. Here, we introduce the OPM-MEG system as a non-invasive investigational tool, with the potential to further detail and explore the structural and functional connectivity of neighbouring cortical areas, with a higher spatio-temporal resolution than currently available. Our initial experiments are consistent with the findings in animal models [33,34] being applicable also to the human brain.

The benefits of OPM-MEG could be important both at research and clinical levels: its higher spatio-temporal resolution would allow to better investigate neural networks, shedding light on the relationships between the connectivity of functionally related brain areas, along with their frequency synchronization. Moreover, this advancement could be applied in clinical populations at different stages, such as those with Alzheimer’s disease. In patients with mild cognitive symptoms, topographical biomarkers based on the analyses of the frequency domain might monitor the progression of the disease over years and help to evaluate therapy response. An even higher impact could be achieved especially at a prodromal (or, even better, preclinical) stage, in which these biomarkers could be used as “gatekeepers” for people at risk of developing Alzheimer’s disease [46].

As this study’s small sample was limited, future research should aim to demonstrate the reproducibility of our results with a larger population. Moreover, it is important to explore the high spatio-temporal resolution of the OPM-MEG system using different stimuli and explore the propagating signals of different brain circuits. Further research is needed to investigate other sensitive pathways in order to better establish the suitability of OPM-MEG for its application in neurophysiological studies. In addition, we have subsequently shown that a factor-six improvement in the DAQ noise floor can be achieved, increasing the SnR of the OPM-MEG for an even higher spatio-temporal resolution.

Based on our observations, OPM-MEG could be a reliable neuroimaging method to identify the activation patterns of close cortical regions in response to specific stimuli. It has the potential to enable reliable neural speed measurements, and spatio-temporal tracking, of propagating signals, including more detailed investigations of the visual pathway.

## ACKNOWLEDGEMENTS

We gratefully acknowledge insightful discussions with Chris Racey and Jamie Ward. This work was supported by the UK Quantum Technologies Hub for Sensors and Timing (EPSRC grant EP/T001046/1) and by the Core Facility ‘Metrology of Ultra-Low Magnetic Fields’ at Physikalisch-Technische Bundesanstalt which receives funding from the Deutsche Forschungsgemeinschaft (DFG KO 5321/3-1 and TR 408/11-1

## AUTHOR CONTRIBUTION

C.A., M.C., T.C., F.D.L, A.G., and P.K. conceived of the presented idea and the data collection techniques, data analysis and interpretation. K.R., T.S., and J.V. oversaw the measurements at PTB, and the design of the visual equipment. All authors discussed the results and contributed to the final manuscript with large inputs from C.A., M.C., T.C., A.G., and P.K.

## DECLARATION OF COMPETING INTERESTS

The authors have no competing interests to declare.

## ADDITIONAL INFORMATION

## Footnotes

‡ https://github.com/OpenBCI/Ultracortex/tree/master/Mark_IV/MarkIV-FINAL

## Notes

### Competing Interest Statement

The authors have declared no competing interest.

